# α-synuclein oligomers displace monomeric α-synuclein from lipid membranes

**DOI:** 10.1101/2023.03.21.533646

**Authors:** Greta Šneiderienė, Magdalena A. Czekalska, Catherine K. Xu, Akhila Jayaram, Georg Krainer, William E. Arter, Quentin Peter, Marta Castellana-Cruz, Kadi L. Saar, Aviad Levin, Thomas Mueller, Sebastian Fiedler, Sean R. A. Devenish, Heike Fiegler, Janet R. Kumita, Tuomas P. J. Knowles

**Affiliations:** Centre for Misfolding Diseases, Yusuf Hamied Department of Chemistry, University of Cambridge, Lensfield Road, Cambridge CB2 1EW, United Kingdom; Cavendish Laboratory, University of Cambridge, J J Thomson Avenue, Cambridge CB3 0HE, United Kingdom; Fluidic Analytics, Unit A, The Paddocks Business Centre, Cherry Hinton Road, Cambridge CB1 8DH, United Kingdom; Department of Pharmacology, University of Cambridge, Tennis Court Road, Cambridge CB2 1PD, United Kingdom

**Keywords:** α-synuclein, oligomers, lipids, aggregation, membranes, Parkinson’s disease

## Abstract

Parkinson’s disease (PD) is an increasingly prevalent and currently incurable neurodegenerative disorder linked to the accumulation of α-synuclein (αS) protein aggregates in the nervous system. While αS binding to membranes in its monomeric state is correlated to its physiological role, αS oligomerisation and subsequent aberrant interactions with lipid bilayers have emerged as key steps in PD-associated neurotoxicity. However, little is known of the mechanisms that govern the interactions of oligomeric αS (OαS) with lipid membranes and the factors that modulate such interactions. This is in large part due to experimental challenges underlying studies of OαS-membrane interactions due to their dynamic and transient nature. Here, we address this challenge by using a suite of microfluidics-based assays that enable in-solution quantification of OαS-membrane interactions. We find that OαS bind more strongly to highly curved, rather than flat, lipid membranes. By comparing the membrane-binding properties of OαS and monomeric αS (MαS), we further demonstrate that OαS bind to membranes with up to 150-fold higher affinity than their monomeric counterparts. Moreover, OαS compete with and displace bound MαS from the membrane surface, suggesting that disruption to the functional binding of MαS to membranes may provide an additional toxicity mechanism in PD. These findings present a unique binding mechanism of oligomers to model membranes, which can potentially be targeted to inhibit the progression of PD.

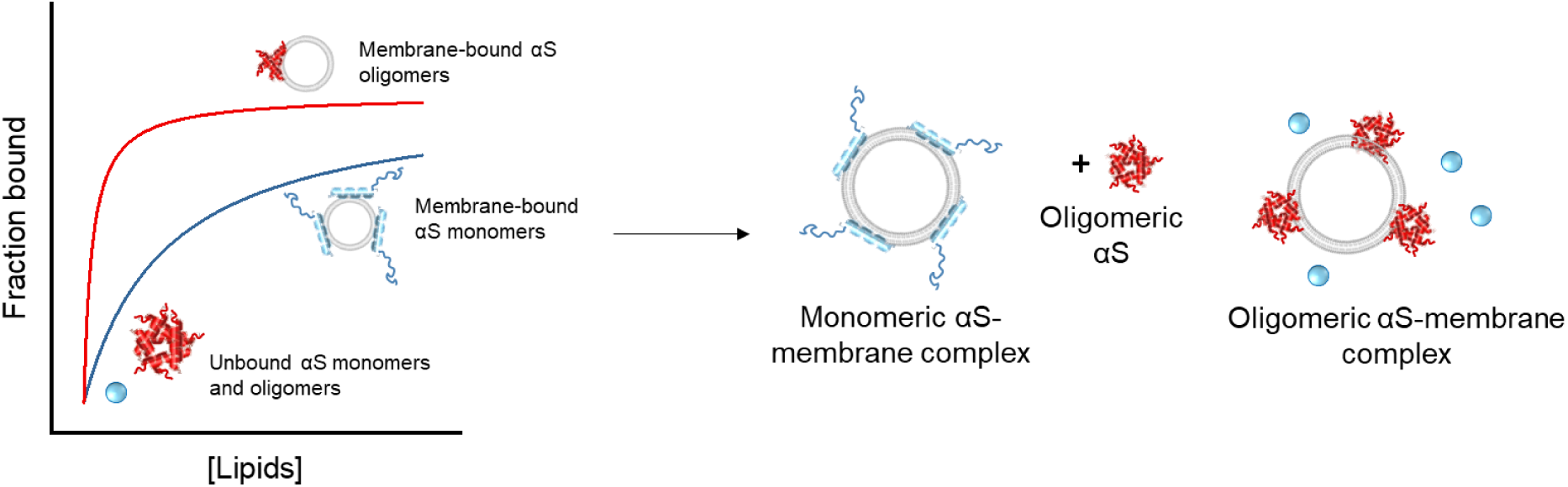

## Introduction

Amyloid fibrils historically constitute the disease-linked proteinaceous inclusions in Parkinson’s disease (PD). These intracellular accumulations, known as Lewy bodies, consist of more than 70 types of molecules, with aggregated α-synuclein (αS) being the main constituent [1]–[3]. However, increasing evidence suggests that oligomeric αS (OαS), rather than fibrils, represent the main neurotoxic agents in PD [4], [5]. The term amyloid oligomers describe low-abundant, transient, heterogenous higher-order protein assemblies that are intermediates in the aggregation pathway [6]–[8]. A range of studies have recently correlated the formation of OαS with cellular toxicity and have shown that oligomer-mediated toxicity can be exerted *via* multiple pathways, including burdening the protein clearance machinery, perturbing the respiratory chain pathway, and importantly, compromising cellular membrane integrity [9]–[14].

The potential role of OαS-membrane interactions in PD has motivated a number of studies focusing on their biophysical characterisation using model phospholipid membranes. OαS have been found to insert into the hydrophobic core of the phospholipid bilayer and disrupt its integrity, thereby leading to leakage of the membrane-surrounded compartments [13]–[15]. By contrast, the binding of monomeric αS (MαS) to such membranes is believed to be crucial for its physiological role in synaptic vesicle trafficking [16]. Several key membrane parameters govern lipid interactions with OαS, including membrane charge, packing density, access to the hydrophobic core, and phospholipid headgroup size [17]–[19]. Accordingly, due to the positively charged N-termini of the polypeptide chains, OαS interact preferentially with negatively charged membranes, composed of phosphatidylserines or phosphatidylglycerols [13], [17]–[19]. Furthermore, low membrane packing density and exposure of the hydrophobic core have been shown to play a key role in toxic OαS-membrane interactions [17]–[19]. Finally, the lipid headgroup size has also been shown to be crucial to OαS association with membranes. OαS preferentially bind to membranes composed of cone-shaped phospholipids with small headgroups, such as phosphatidylglycerol and cardiolipin [19].

While the binding of MαS to model lipid bilayers *in vitro* is well-described quantitatively, based on data obtained from circular dichroism (CD) spectroscopy and diffusion times of particles (fluorescence correlation spectroscopy (FCS) and microfluidic diffusional sizing (MDS)) [20]–[24], thermodynamic quantification and mechanistic studies of oligomer binding to lipid membranes are still challenging to perform. This is largely due to the structural heterogeneity, transient nature, and low abundance of oligomeric species [14]. Previous attempts to quantify OαS-membrane binding affinity have been limited to surface plasmon resonance (SPR) studies using flat membranes and non-purified oligomer samples, which may contain fibres, potentially distorting the oligomer-membrane binding affinity values obtained [25], [26]. Alternative strategies include the fluorescence quenching assay, in which the fluorescence arising from tryptophan or a lipid-specific dye incorporated into large unilamellar vesicles (LUVs) is reduced upon membrane binding of OαS and HypF-N oligomers [27], [28]. Moreover, the majority of the optical microscopy studies of OαS-membrane interactions are carried out with giant unilamellar vesicles (GUVs), which do not provide an accurate representation of the highly curved synaptic vesicles found in the presynaptic termini, the primary cellular location of MαS [18], [29], [30]. This limitation is particularly significant, as the curvature of the membrane is known to affect the binding activity of MαS and OαS to the membrane [20]. Rapid and robust methods that enable the quantification of OαS-membrane interactions in native-like conditions are therefore required.

Here, we provide quantitative insights into the binding of OαS with model liposome membranes using an array of microfluidics-based assays that enable in-solution quantification of OαS-membrane interactions. Specifically, we utilize microfluidic diffusional sizing (MDS) and microfluidic free-flow electrophoresis (μFFE) to gain an understanding of the mechanisms which lead to neurotoxic effects in PD. We find that both MαS and OαS lower the absolute negative charge of liposomes upon their binding, reducing the ζ-potential of the complex, thus enabling a decreased liposome stability and increased vesicle merging propensity. We further demonstrate that OαS bind to membranes with up to a 150-fold higher affinity than their monomeric counterparts. Moreover, we show that OαS compete with and displace MαS from the membrane surface, potentially disrupting functional binding of MαS to membrane surfaces. Taken together, our study reveals important mechanistic details of MαS and OαS membrane binding interactions and presents a new route of toxicity in PD, which involves MαS loss of function.

## Results

### Probing OαS and MαS binding to the lipid membranes

In cells, MαS is primarily located at the synaptic termini close to synaptic vesicles and the inner plasma membrane leaflet, both of which vary in chemical composition and physicochemical membrane properties. Phosphoserine and phosphocholine lipids are the main negatively charged and zwitterionic phospholipid species present in synaptic vesicles and the inner plasma membrane leaflet, respectively [31], [32]. To build mechanistic understanding of MαS/OαS-lipid interactions involvement in Parkinson’s disease, we therefore explored the interactions between MαS/OαS with a simple model membrane, composed of 1,2-dioleoyl-sn-glycero-3-phospho-L-serine (DOPS) and 1,2-dioleoyl-sn-glycero-3-phosphocholine (DOPC) [30]. Both lipids form bilayers in the liquid crystalline phase under room temperature conditions due to the presence of unsaturated acyl chains. These unsaturated acyl chains facilitate MαS-membrane binding and its related oligomerisation propensity and are thus particularly relevant model systems in αS-linked PD pathology [24], [33], [34].

OαS is a broad term referring to heterogenous higher-order protein assemblies [8]. In our study, we chose to work with *in vitro* generated, chemically unmodified and well-characterised oligomeric αS species of a known structure [14]. The isolated oligomeric species consist of 30–40 monomeric subunits with an *R*_h_ of 10.1 ± 1.0 nm (Figure S1a, b) as determined by analytical ultracentrifugation and MDS respectively. The isolated OαS are kinetically trapped species on the aggregation pathway and remain stable for 2 days, after which they dissociate back into MαS (Figure S1c). Generally speaking, OαS possess different membrane permeabilization propensities, ranging from toxic, β-sheet rich membrane active species to amorphous aggregates that do not compromise membrane integrity [8], [13], [26], [35]. Our *in vitro* generated OαS do not compromise DOPS membrane integrity over 5 hours as indicated by the results of the calcein membrane leakage assay (Figure S1d).

To probe OαS binding to liposomes, we first performed CD spectroscopy experiments, a well-established method for monitoring the secondary structure changes of MαS upon membrane binding [23]. While MαS undergoes a structural transition from intrinsically disordered to an α-helix-rich fold upon incubation with negatively charged, monosaturated DOPS LUVs, as evident by the minima at 209 nm and 223 nm in the CD spectra (Figure S2a) [16], [36], [37], OαS do not undergo significant structural changes upon incubation with DOPS LUVs (Figure S2b) [13]. The minimal changes observed by CD spectroscopy can be due to a lack of interaction between the oligomers and lipids, or to a lack of structural change in the oligomers upon binding.

### Validation of the MDS assay for quantification of MαS- and OαS-lipid interactions

As an alternative method to probe OαS-vesicle complex formation, we turned to MDS, which does not rely on changes in secondary structure. MDS exploits differences in diffusive mass transport in a laminar flow regime to characterise bound and unbound molecules based on their hydrodynamic radii (*R*_h_) [22], [38]–[41].This is achieved by co-flowing the sample with a buffer in a microfluidic channel (Figure 1a). Under laminar flow conditions, the analyte mixes with the buffer by diffusion only. The rate of diffusion and hence the profiles of the diffused analytes depend on the size of the particles and thus can be used to determine the hydrodynamic radius of the analyte, and thereby the degree of binding [22].

**Figure 1.**
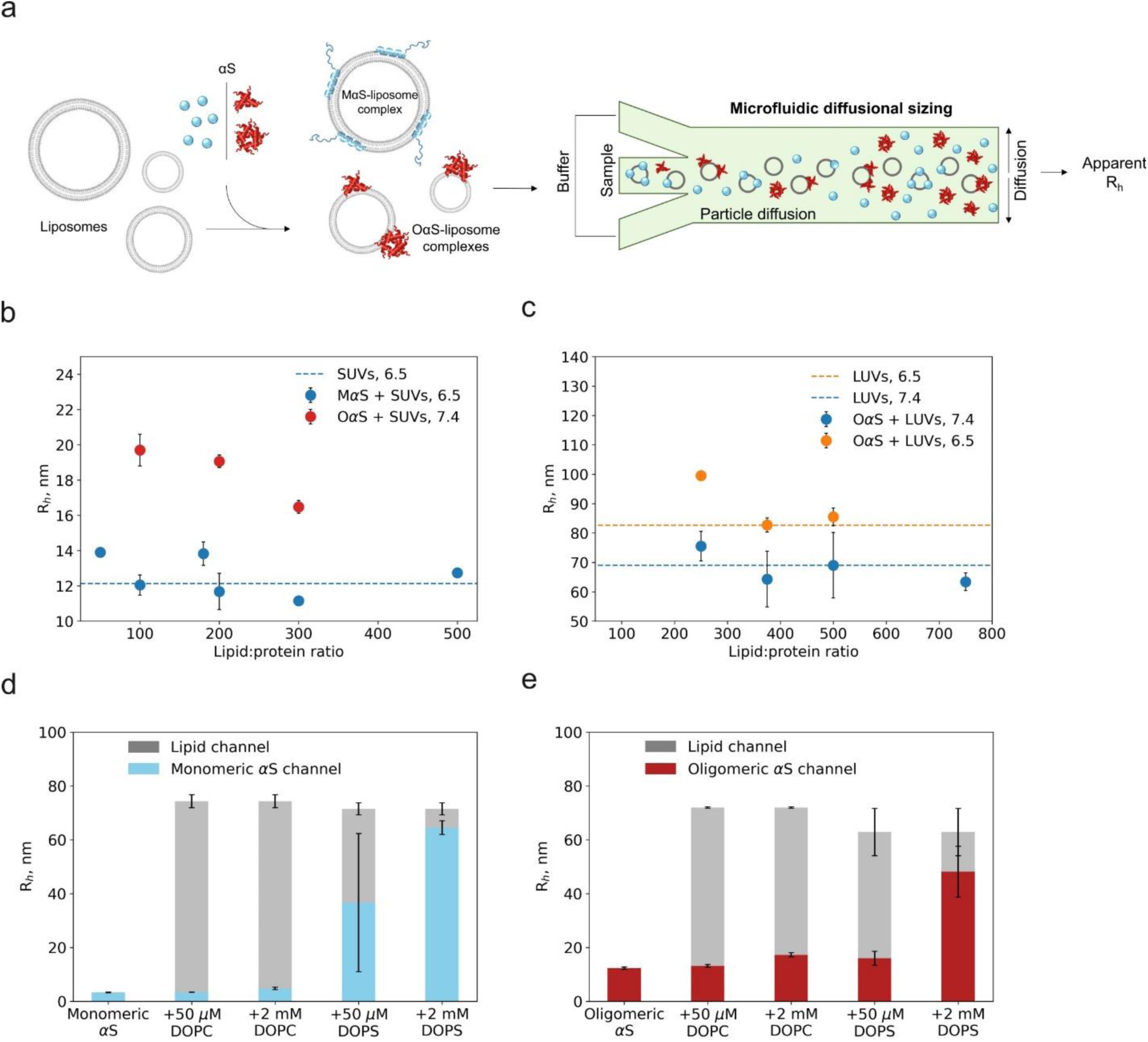
Probing MαS- and OαS-membrane interactions. (a) Microfluidic diffusional sizing-based approach for probing MαS- and OαS-membrane interactions. Alexa 488 labelled MαS or OαS were mixed with two different concentrations of ATTO 647-labelled DOPS or DOPC vesicles. The equilibrated mixtures were injected into an MDS chip. A conventional widefield fluorescence microscope was used to record fluorescence intensity of the samples excited with 488 nm and 647 nm light simultaneously, allowing us to probe protein, protein-lipid complex and lipid vesicle sizes in the same sample. Vesicle size measurements of (b) MαS/OαS-SUV and (c) OαS-LUV mixtures. Proteins were mixed with liposomes at different rations, then sized by MDS. Data represent the hydrodynamic radii measured in the lipid channel. Dashed lines indicate sizes of pure liposomes. MDS data on binding of MαS (d) and OαS (e) to zwitterionic (DOPC) and negatively charged (DOPS) LUVs at pH 6.5. Two lipid concentrations were used – low sub-saturating (50 µM) and high saturating (2 mM). Hydrodynamic radii (*R*_h_, nm) measured from both protein (MαS - blue, OαS - red) and lipid channel (grey) are represented with bars. 2 μM MαS and OαS (monomer equivalents) in 20 mM NaP pH 6.5 buffer was used. Error bars represent standard deviations of n = 3–4 measurements on individual microfluidic chips.

The MDS binding affinity measurements rely on protein size increase upon addition of binding partners (liposomes). This is then converted to the liposome-bound fraction of the protein, assuming that the radius of free unbound protein is equal to 0 and the radius of fully bound protein in a liposome-protein complex is equal to 1. Hence it is critical to ensure that measured changes in diffusive properties and thereby sizes reflect complex formation rather than other events, such as lipid membrane deformation by MαS [42]. To test the hypothesis, we measured liposome sizes in the presence of varying MαS/OαS concentrations by MDS (Figure 1b, c). We found that vesicle size remains constant at different lipid:protein ratios in MαS-SUV and OαS-LUV samples (Figure 1b and c, respectively). Importantly, sizes of OαS-LUV and MαS-SUV complexes are comparable to the sizes of pure SUVs and LUVs (indicated by blue dashed line in Figure 1b, orange and blue dashed lines in Figure 1c). This indicates that the sizes of liposomes remain unchanged upon binding of MαS and OαS over a large range of lipid:protein ratios tested and suggests that the diffusive properties of MαS-SUV, OαS-LUV complexes and pure DOPS SUVs/LUVs remain the same across different experimental conditions. This experimental validation indicates that the measured average *R*_h_ values are a good proxy to determination of the liposome bound fraction of MαS and OαS and thereby binding affinity.

Taken together, we have validated the MDS assay for quantifying MαS-SUV, MαS-LUV and OαS-LUV interactions and confirmed that it is indeed suitable for our study goals.

### OαS selectively bind to negatively charged lipid membranes

As a first step, we probed MαS binding to LUVs (*R*_h_ = 74.3 ± 2.4 nm). The *R*_h_ of pure MαS was measured to be 2.6 ± 0.6 nm, in agreement with previous measurements (Figure 1d) [22]. No MαS interactions with DOPC LUVs were detected, indicated by the hydrodynamic radius remaining constant across the pure MαS and MαS + DOPC samples (Figure 1d). By contrast, in the presence of negatively charged DOPS LUVs, MαS (2 µM) forms a complex with DOPS LUVs, as indicated by the change of *R*_h_ from 12.3 ± 0.45 nm to 48.1 ± 9.4 nm at 2 mM DOPS concentration (Figure 1d). At 50 μM DOPS concentration, the measured size is 36.7 ± 25.7 nm, indicating that only a fraction of MαS is bound to the DOPS vesicles with the rest remaining in the unbound state in solution, hence the large standard deviation (Figure 1d). This further confirms that MαS-lipid interactions are charge-specific, and a negative membrane charge is required for MαS binding [16].

Next, we probed OαS-membrane interactions using MDS and successfully applied the method for quantifying OαS-membrane binding. The measured *R*_h_ of pure OαS is 12.3 ± 0.5 nm, notably larger than that of MαS (Figure 1e). OαS do not form complexes with zwitterionic DOPC LUVs, as indicated by the *R*_h_ of OαS remaining constant after the addition of lipids at 50 μM and 2 mM (Figure 1e). By contrast, as in the case of MαS, OαS forms a complex with the negatively charged DOPS LUVs, as indicated by the increase of the apparent radius to 48.1 ± 9.5 nm at 2 mM DOPS concentration (Figure 1e). These results therefore demonstrate that oligomers are able to bind DOPS LUVs, but with minimal rearrangement in secondary structure, making such interactions particularly challenging to detect using CD spectroscopy.

Taken together, both MαS and OαS preferentially bind to negatively charged membranes, *via* interactions governed largely by electrostatic forces, in accordance with previous work [16], [19]. This commonality in charge binding specificity suggests that regions mediating both MαS- and OαS-membrane interactions overlap [16], [43], [44].

### OαS bind more tightly to highly curved membranes

In biological settings, MαS is exposed to membranes with variable curvatures, namely highly curved synaptic vesicles of 30–40 nm diameter and plasma membrane, which can be considered flat on the scale of MαS. However, very little is known about how membrane curvature affects OαS binding. To investigate this, both MαS and OαS were incubated with DOPS SUVs and LUVs, based on the results presented in Figure 1c, to mimic the curvatures of synaptic vesicles and the plasma membrane surface, respectively. The sizes of the protein-lipid complexes were then measured using MDS.

When MαS and OαS are mixed with DOPS SUVs (*R*_h_ = 14–15 nm), the particle sizes measured is comparable to that of lipid vesicles at both low (50 μM) and high (2 mM) lipid concentrations and at both pH 6.5 and 7.4 (Figure 2a, b), indicating that all protein is recruited to the membrane surface under these conditions. By contrast, in when MαS and OαS were mixed with 50 μM DOPS LUVs (*R*_h_ = 66–73 nm), the particle sizes measured in the protein channel were intermediate between the sizes of free MαS/OαS and the LUVs, indicating that only a fraction of MαS/OαS formed complexes with the flatter membrane surface of DOPS LUVs compared to SUVs (Figure 2c, d).

**Figure 2.**
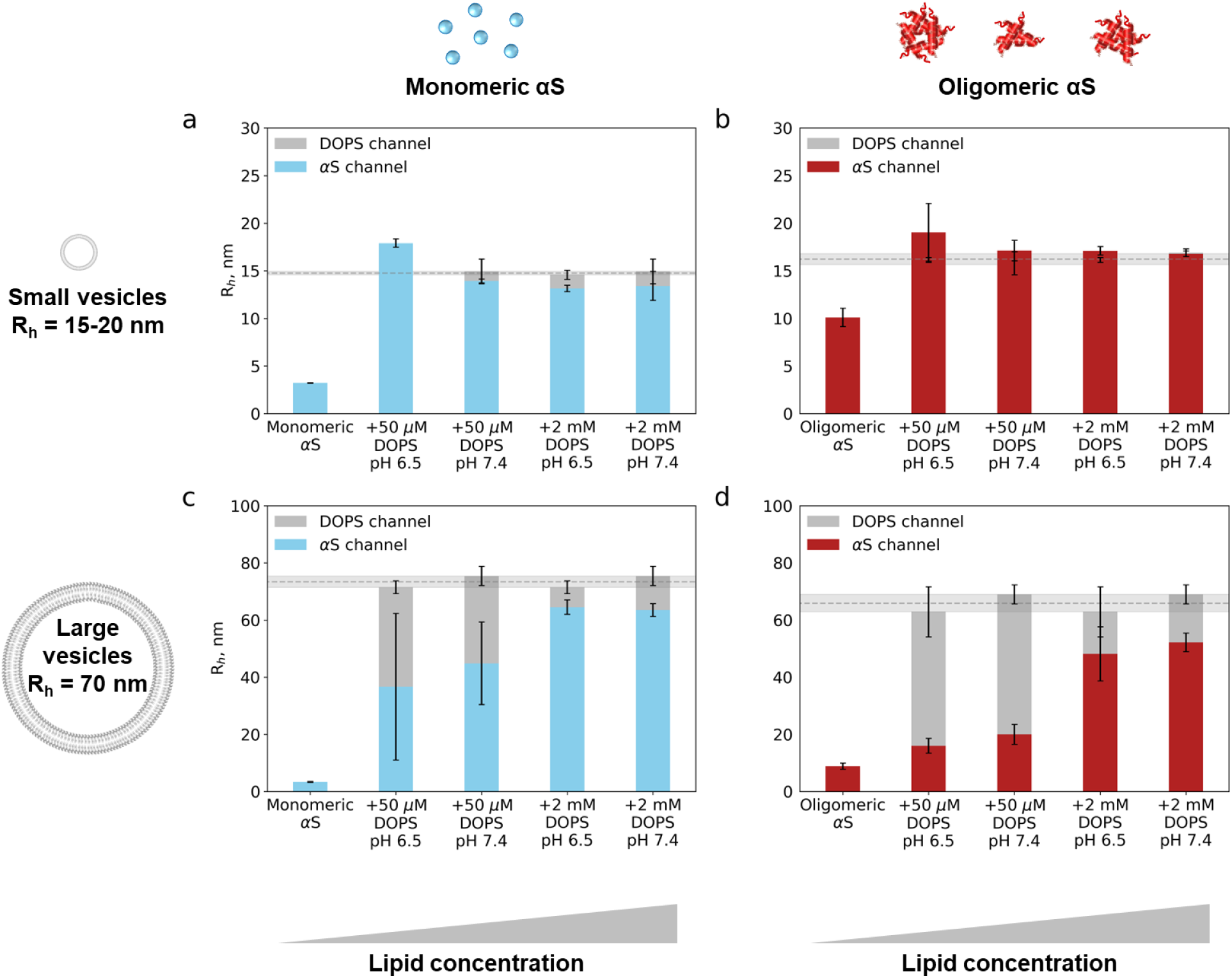
Binding of MαS (a, c) and OαS (b, d) to DOPS SUVs and LUVs. Two lipid concentrations were used – low, sub-saturating (50 µM) and high, saturating (2 mM). Hydrodynamic radii (*R*_h_, nm) measured both in the protein (MαS - blue bars, OαS – red bars) and the lipid channel (grey bars and grey lines). 2 μM MαS and OαS (monomer equivalents) in 20 mM sodium phosphate buffer was used. Error bars represent standard deviations of n = 3–4 measurements on individual microfluidic chips.

Taken together, these results suggest that membrane curvature can affect both OαS and MαS membrane binding. Both MαS and OαS bind more tightly to highly curved SUVs rather than LUVs, as previously shown [45], indicating that both MαS and OαS-membrane interactions may be governed by similar physicochemical mechanisms.

### OαS have a higher membrane binding affinity than MαS

To better understand the OαS structure-membrane binding relationship, we turned to quantitative characterisation and comparison of MαS- and OαS-membrane binding by the means of MDS We measured the binding affinity parameters of MαS-SUV, OαS-LUV, OαS-SUV complexes assuming a two state model (unbound and membrane bound αS).The binding curves were recorded by measuring the radii under conditions, where MαS and OαS are in the free unbound, partially vesicle-bound and fully vesicle-bound states respectively, after samples reached the equilibrium state. The diffusion profiles in both protein and lipid fluorescence channels were recorded simultaneously; the radii from the protein channel report on the degree of protein binding to the liposomes (Figure 3), whilst the radii from the lipid channel serve as a control confirming that sizes of the liposomes remain unaffected upon MαS/OαS binding (Figure 1b, c). The raw radii from the protein channel are then normalized for each sample individually to determine the liposome-bound MαS/OαS fraction. We have found that OαS bound to the DOPS SUVs more strongly than MαS at both pH 6.5 and 7.4, evidenced by their decreased *K*_D_ values to low nanomolar level (Figure 3a, b, Table S1). As the calculated *K*_D_ values reported here are expressed in units of molar monomer particle-vesicle interactions, this higher affinity for the OαS indicates that OαS bind more strongly to vesicles than MαS. Moreover, the effective stoichiometry (expressed as lipid molecules per binding site) of lipids per OαS particle increased by more than an order of magnitude, indicating that fewer OαS can bind per vesicle as a result of steric occlusion or membrane remodelling by OαS.

**Figure 3.**
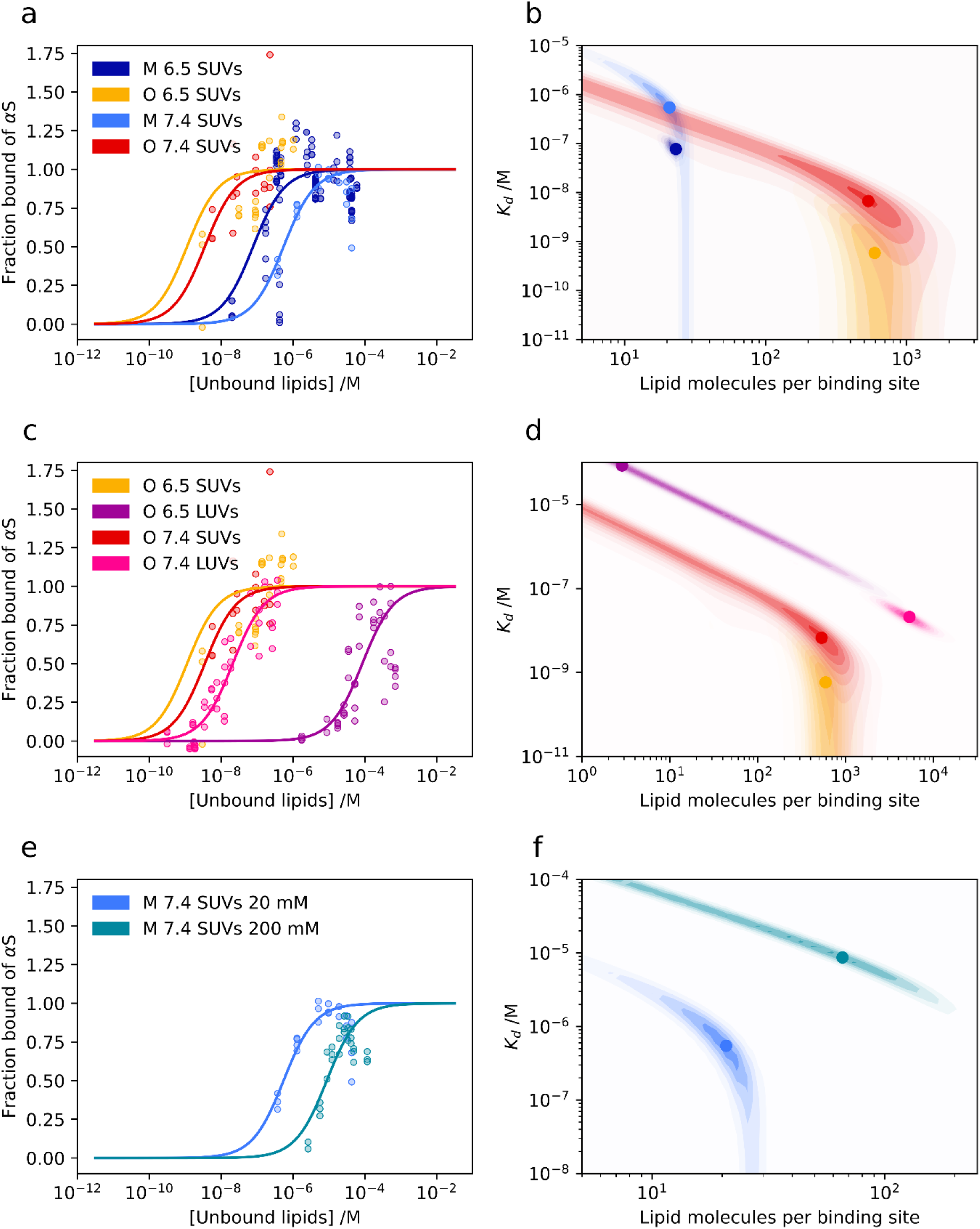
Quantification of MαS/OαS binding affinities and stoichiometries to negatively charged DOPS vesicles with the MDS. Data (points) and fits (lines) are shown in the left panels, alongside the corresponding marginal posterior probability distributions over affinity (*K*_D_) and stoichiometry expressed as lipid molecules per binding site in the right panels. (a, b) Binding of both MαS and OαS to SUVs at both pH 6.5 and 7.4. (c, d) Binding of OαS to SUVs and LUVs at pH 6.5 and 7.4. (e, f) Binding of MαS to SUVs at pH 7.4, with varying concentrations of sodium phosphate buffer (20 and 200 mM).

### Effect of pH change on OαS-membrane binding is membrane curvature-dependent

MαS is exposed to a wide range of pH environments in the cell. While in the neuronal synapse, which is the primary location of αS, the pH is 7.4 [46], misfolded forms of MαS can be found in lysosomes under pH 6.5 conditions, where they appear after internalisation during disease spreading [46]–[48]. Furthermore, MαS has been shown to interact with a variety of different membranes including synaptic vesicles and cellular plasma membrane with different binding strengths at different pH levels [49]. To better elucidate the binding strength of OαS and MαS under a range of physiological conditions, we measured the binding affinities between OαS and DOPS SUVs at pH 6.5 and 7.4 by the means of MDS.

We found that the change in pH only had a minor effect on MαS/OαS-DOPS SUVs binding parameters (Figure 3a, b). For both MαS and OαS, the membrane binding affinity is less than one order of magnitude higher at pH 6.5 than pH 7.4 conditions (Figure 3a, b). In both cases, the binding stoichiometry is also unaffected by the pH (Figure 3a, b). The slightly higher binding affinity at pH 6.5 may, however, result from reduced electrostatic repulsion between the negatively charged DOPS headgroups and the negatively charged C-terminus of αS [16].

By contrast, the environmental pH has a clear effect on the binding of OαS to DOPS LUVs (Figure 3c, d). At pH 7.4, OαS bind to LUVs with a markedly higher affinity and stoichiometry than at pH 6.5. The drastic effect of the pH change on the binding parameters between OαS and relatively flat LUVs, but not on highly curved SUVs, suggests that membrane curvature rather than electrostatic interactions determine the strength of OαS-DOPS vesicle interactions. In addition to the differential effects of pH on the OαS-SUV and -LUV binding systems, we observed that the affinity of OαS for SUVs is higher compared to LUVs. These results indicate that the membrane curvature-sensing ability is not lost upon oligomer formation, consistent with the notion that the same residues are involved in membrane binding in both the MαS and OαS [17], [19].

### MαS-membrane interactions are electrostatically driven

Further investigation revealed that MαS binds to SUVs with a two-fold lower binding affinity at high ionic strength conditions (200 mM NaP c.f. 20 mM NaP for all other binding experiments) (Figure 3e, f and Table S1). The ratio of the *K*_D_ and binding stoichiometry is well-constrained, differing by ∼2 times between the two salt concentrations. This is in good agreement with previous studies, which demonstrate the importance of ionic interactions in MαS-membrane binding that can be inhibited by high concentrations of ions [50].

Taken together, while OαS interact with the negatively charged DOPS membranes more tightly than MαS OαS interaction with the membranes of varying curvature and different pH levels did not yield a definitive trend. This further demonstrates the mechanistic variations of OαS- and MαS-membrane interaction, affected by membrane curvature and pH.

### MαS forms a single type of complex with DOPS liposomes

To further explore the nature of interactions between MαS or OαS and DOPS vesicles, we utilised microfluidic free-flow electrophoresis (μFFE) to probe αS-membrane interactions in solution in higher detail. μFFE electrophoretically separates fluorescently labelled species lateral to their direction of flow and thus facilitates the determination of their electrophoretic potential based on their deflection profiles (Figure S4). Using μFFE, we were able to separate complex protein-lipid mixtures on-chip and thereby determine their ζ-potential, which is an indicator of particle stability (Figure 4a) [51].

**Figure 4.**
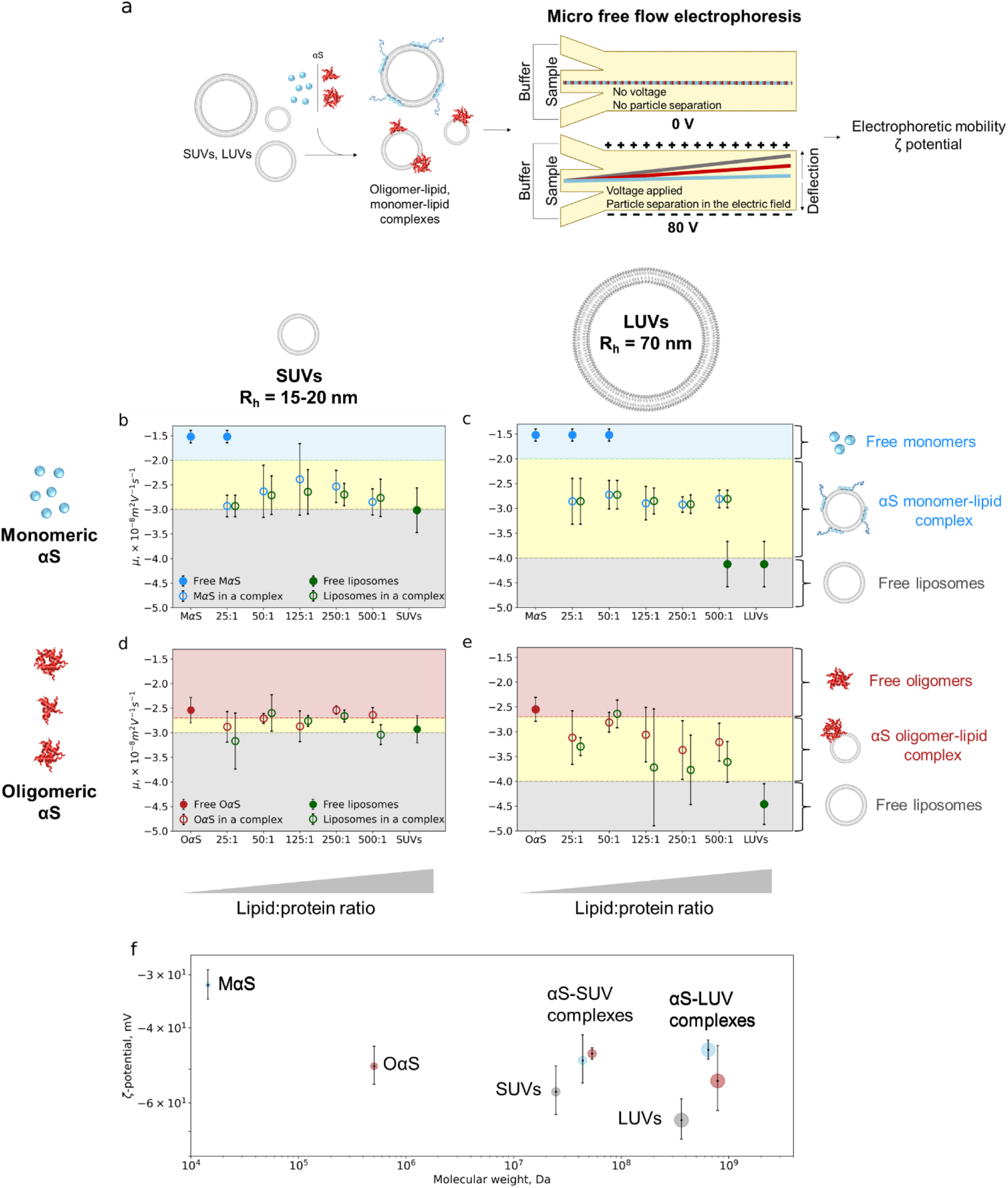
Electrophoretic analysis of MαS/OαS-lipid complexes by micro free flow electrophoresis (μFFE). (a) Experimental scheme of μFFE. Electrophoretic mobilities (µ, × 10^-8^ m^2^ V^−1^ s^−1^) of αS-liposome complexes: (b) MαS with SUVs, (c) MαS with LUVs, (d) OαS with SUVs, € OαS with LUVs. In each plot, first the mobility of free protein (full blue/red circles) is plotted, followed by the mobility values at different molar lipid:protein ratio (25:1, 50:1, 125:1, 250:1, 500:1; empty circles) with the mobility of the free SUV/LUV plotted last (full green circles). Blue, red, yellow and grey regions represent the mobility ranges of MαS, OαS, protein-lipid complexes and free vesicles respectively. ζ-potentials (f) of MαS/OαS and MαS/OαS-liposome complexes, determined through the combination of the μFFE and MDS measurements (plotted as the function of molecular weight (Da)). The marker sizes are drawn to a relative scale of MαS, OαS, MαS/OαS-liposome complexes and pure liposome dimensions. All experiments were run in 20 mM pH 6.5 NaP buffer. Each data point represents the mean of n>3 independent repeats, and error bars represent standard deviations. The blue, red, yellow and grey regions are guide to the eye and represent mobilities of free MαS, OαS, MαS/OαS-liposome complexes and pure liposomes.

First, we investigated MαS interactions with both DOPS SUVs and LUVs (Figure 4b, c). The measured electrophoretic mobility of MαS was -1.52 × 10^−8^ m^2^ V^−1^ s^−1^, in good agreement with the literature data [52]. Upon increasing the concentration of liposomes, we observed that MαS is recruited to the liposomes, as indicated by the presence of 2 separate species at low lipid-to-protein (L:P) ratios (Figure 4b, c). Notably, the mobility of the MαS-lipid complex does not change across different L:P ratios (ca. -2.68 × 10^−8^ m^2^ V^−1^ s^−1^ and ca. -2.85 × 10^−8^ m^2^ V^−1^ s^−1^ for LUVs and SUVs complexed with MαS, respectively). The electrophoretic mobility is determined by the hydrodynamic radius and charge of the species, so its constant value at different L:P ratios suggests that the composition of the MαS-lipid complex is not dependent on the L:P ratio. The presence of these two well-defined species indicates that MαS forms a single type of complex of the same electrophoretic mobility with DOPS liposomes at all L:P ratios (Figure 4b, c).

### OαS protein-membrane complexes vary based on lipid concentration

Next, we investigated OαS interactions with both SUVs and LUVs (Figure 4d, e). The mobility of OαS is two times higher compared to the mobility of MαS (–2.54 ± 0.26 × 10^−8^ m^2^ V^−1^ s^−1^ vs -1.52 ± 0.12 × 10^−8^ m^2^ V^−1^ s^−1^), due to their higher charge arising from their multiple monomer subunits, and corresponding sub-linear increase in hydrodynamic radius [52]. Interestingly, OαS show different electrophoretic mobility patterns when mixed with DOPS liposomes than that of MαS. At a low L:P ratio, all OαS are bound to the DOPS liposomes, as indicated by a single high-mobility species detected in the protein channel (Figure 4d, e). This is attributable to a higher DOPS membrane binding affinity and a smaller number of oligomers present in the OαS-lipid mixture at the same L:P ratio, since the protein concentration is given in monomer equivalents. Moreover, the mobility of the OαS-DOPS complex increases with increasing lipid concentration and is less constrained in comparison to the constant mobility of the MαS-lipid complexes, indicating that different complexes, likely of different bound oligomer:lipid stoichiometry, form at different L:P ratios (Figure 4d, e). This difference between the behaviours of MαS and OαS could be due to the observed steric occlusion effects of OαS, which occupy a larger area of the vesicle surface [53].

### MαS and OαS reduce the effective surface charge of DOPS liposomes

Another parameter that is especially relevant for charged particles in aqueous solutions is the ζ-potential, which can be calculated from measured electrophoretic mobilities and hydrodynamic radii. The ζ-potential describes the electrical potential at the edge of the interfacial double layer of ions and counterions near the surface of a charged particle. We thus calculated the ζ-potentials of MαS, OαS, DOPS SUVs and LUVs, and the protein-lipid complexes formed at saturating lipid concentrations (Figure 4f). The calculated ζ-potentials of MαS/OαS-DOPS complexes range between -30–70 mV, indicating moderate stability of the complexes (Figure 4f) [54], [55].

Notably, the ζ-potentials of the MαS/OαS-liposome complexes are lower than those of the liposomes alone (Figure 4f), suggesting that the negative DOPS liposome surface charge is screened by the membrane surface-bound MαS and OαS. The reduced ζ-potential of the MαS/OαS-liposome complexes also suggests that the MαS/OαS-liposome complexes may have different stability properties compared to those of pure liposomes.

### OαS displace MαS from the lipid membranes

MαS and OαS coexist in the complex cellular environment and can simultaneously interact with a wide range of binding partners, including lipid membranes [49], [56]. Given the similarities between the binding of OαS and MαS to vesicles, we hypothesised that the two species can compete for binding sites on a membrane surface. We thus carried out a competition assay between OαS and MαS with MDS. MαS, OαS, and DOPS lipid vesicles were labelled with Alexa 546, Alexa 488, and DOPE-ATTO 647, respectively, to monitor the formation and composition of MαS/OαS-lipid complexes. MαS (2 μM) was first incubated with a sub-saturating concentration of DOPS lipids (150 μM), followed by the addition of 2 μM OαS. The mixture was probed by MDS both before and after incubation with oligomers, with all three components monitored separately by their orthogonal fluorescent labels (Figure 5).

**Figure 5.**
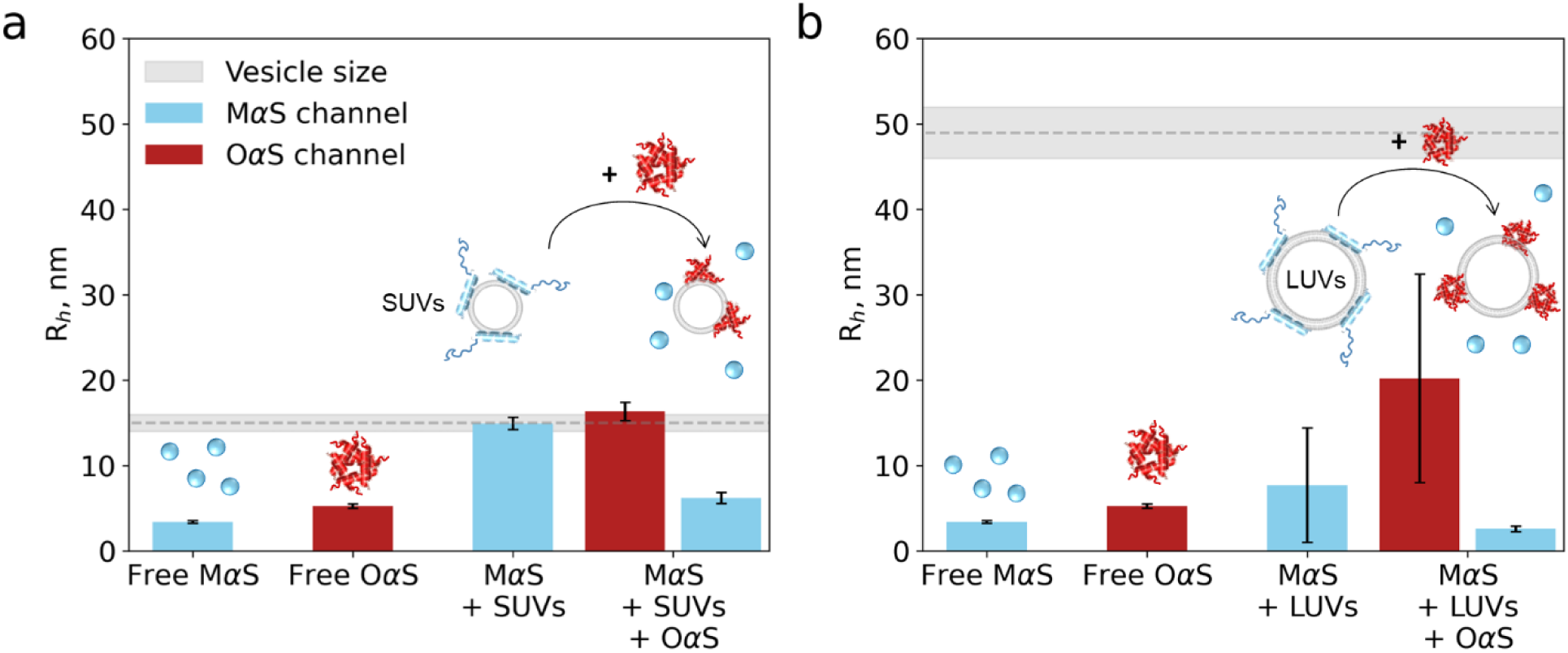
OαS-MαS binding competition assay. MαS (2 μM) was mixed with 150 μM SUVs (a, *R*_h_ = 15 ± 1 nm) and LUVs (b, *R*_h_ = 49 ± 3 nm, shown in grey) and allowed to equilibrate, following which the size was measured by the MDS. Following this, OαS (2 μM) were added and further left for equilibration. First, sizes of free protein are depicted for comparison (MαS: *R*_h_ = 3.4 ± 0.1 nm, OαS: *R*_h_ = 5.2 ± 0.2 nm). Then, a size measured from the MαS channel after incubation with lipid vesicles is plotted. (*R*_h_ = 14.9 ± 0.7 nm for SUVs and *R*_h_ = 7.7 ± 6.7 nm for LUVs). Upon addition of OαS however, the size of MαS decreases (*R*_h_ = 6.2 ± 0.7 nm for SUVs and *R*_h_ = 2.6 ± 0.3 nm for LUVs), while the size measured for the OαS channel is larger (*R*_h_ = 16.3 ± 1.1 nm for SUVs and *R*_h_ = 20.2 ± 12.2 nm for LUVs) in comparison to unbound protein (*R*_h_ = 5.3 ± 0.2 nm). Error bars represent standard deviations of n = 3–4 measurements on individual microfluidic chips.

As shown in Figure 5a, the apparent size of MαS increased from *R*_h_ = 3.4 ± 0.1 nm to 14.9 ± 0.7 nm following the addition of DOPS SUVs (blue bars), indicating the formation of the MαS-lipid complex (Figure 5a). Following the addition of OαS, the MαS complex size decreased to *R*_h_ = 6.2 ± 0.7 nm, indicating that a fraction of the lipid-bound monomer dissociated back into the solution from the membrane surface (Figure 5a). By contrast, the size of OαS increased from *R*_h_ = 5.3 ± 0.2 nm to 16.3 ± 1.1 nm following their incubation with the equilibrated MαS-SUV mixture, indicating that OαS was able to displace the bound MαS and bind to the SUVs (red bars in Figure 5a). The same trend was observed in the MαS-OαS-LUV system (Figure 5b). First, MαS binds to DOPS LUVs, as indicated by a size increase from *R*_h_ = 3.4 ± 0.1 nm to 7.7 ± 6.7 nm (blue bars), and then is outcompeted by OαS, as indicated by a decrease in size to *R*_h_ = 2.6 ± 0.3 nm. The dissociation of MαS is a direct consequence of OαS binding to the membrane as MαS-vesicle samples incubated for longer than 1 hour did not exhibit any further dissociation of MαS (Figure S5).

To rule out the possibility of the dissociation of MαS from OαS during the experiments, we co-incubated MαS and OαS for 2 hours and analysed the mixtures by MDS (Figure S6). Following incubation, the sizes of MαS and OαS remained unchanged (*R*_h_ = 2.1 ± 0.8 nm and 12.3 ± 0.1 nm, respectively) from the measurements at t = 0 h (*R*_h_ = 2.5 ± 0.8 nm and 10.0 ± 1.4 nm, respectively). The stable size of OαS throughout the experimental timeframe indicates that there is neither growth nor dissociation of OαS when incubated with MαS.

We thus suggest that when membrane surface area is limited, OαS can displace MαS from the membrane surface. This demonstrates the reversibility of MαS-membrane binding, which can be modulated by the presence of OαS; OαS, as a stronger binder, is able to out-compete MαS in binding to vesicles in solution.

## Discussion

While OαS have been shown to induce toxicity through aberrant interactions with cellular membranes [13], additional fundamental biophysical insights into the parameters governing their interactions with lipid bilayers promoting OαS pathological function are required. In this study, we addressed this challenge and quantitatively described the interactions of MαS and OαS with lipid bilayers through capitalising on an array of solution-phase microfluidics approaches. Such in-solution measurements characterised by short experimental timescales and low sample consumption allowed us to quantitatively characterise, for the first time, the nature and physicochemical parameters defining the binding of OαS to lipid bilayer membranes.

Such detailed studies have previously remained challenging, due to intricacies both in the preparation of OαS systems, and in the availability of suitable quantitative characterisation methods. Yet, using the microfluidic approaches described herein, we have been able to determine the binding mechanism and affinity of OαS–membrane interactions. By employing two commonly used model lipid systems, we determined that MαS bind to the negatively charged DOPS membranes with mid-nanomolar to low-micromolar affinity, in good agreement with previous studies [20]–[22], while OαS binds to DOPS membranes with low nanomolar affinities, with the *K*_D_ values being in the same range as previously shown for heterogenous protofibrillar αS [25], [26]. This finding is of particular importance, as binding affinity data of aggregated αS have only been available for non-purified, heterogenous protofibrillar αS samples, likely containing fibres and MαS, whilst we measured membrane binding affinities of pure OαS [25].

We further explored the effect of pH on αS–membrane binding affinity, as protein pH charge dependence suggests that it can regulate the degree of αS membrane association. At two pH levels, 6.5 and 7.4, the charge of MαS differs by 1 charge unit (theoretical values at pH 7.4 and 6.5, are -9.1 and -8.3 respectively) [57]. However, the membrane binding affinities of MαS and OαS are similar at both pH levels tested (Figure 5). This suggests that the membrane binding ability of αS is not affected by mild pH changes in the cellular environment, thus maintaining its crucial role in synaptic vesicle trafficking and other αS-assisted membrane remodelling events [58].

Notably, we find that OαS-membrane interactions are up to 150-fold stronger than those of MαS binding to the negatively charged lipid bilayer. This dramatic increase in binding strength could be attributed to the high avidity of the OαS particles. OαS can expose multiple N-termini that are required for OαS anchoring to the membrane, thus increasing the overall binding strength [59]. A similar effect has been observed for transthyretin, which exhibits a higher membrane binding propensity in an aggregated form [60].

The presented detailed analyses of MαS- and OαS-lipid interactions highlight the key role of electrostatic forces in governing OαS–membrane interactions. We have thus demonstrated that a negative membrane charge is required for binding, which is in accordance with previous reports for MαS and OαS [17], [19], [20], [30]. However, cellular membranes typically consist of mixtures of lipids, including PS/PC/PE and others [32], [61]. Previous studies have indicated that the binding affinity of MαS to membranes decreases substantially, up to 10^3^ times, with increasing PC content [20], [21]. Our results, along with previous reports, highlights the significant role of electrostatic interactions in governing OαS-membrane binding strength [19]. Based on this understanding, we hypothesize that the binding affinity of OαS to complex membranes would similarly decrease with increasing PC/PE content.

Membrane curvature is another crucial factor in governing αS-membrane binding. Preferential binding of MαS to SUVs in comparison to LUVs manifested with 100-fold differences in affinity is generally attributed to the curvature match between SUVs and that of the αS α-helix, thus favouring the interaction and increasing αS-membrane binding affinity [20], [21], [62]. The increased lipid headgroup area of SUVs compared to LUVs, which is directly linked to the exposure of the membrane hydrophobic core, does not seem to play a role in MαS-membrane binding affinity, as MαS adsorbs on the membrane surface and does not deeply penetrate into the hydrophobic core of the bilayer [63]. We observe a similar trend for OαS, where binding affinity increases by an order of magnitude as the size of the vesicles decreases. This can be due to differences in membrane packing and exposure of the hydrophobic core in membranes with higher surface curvature [13], [14], [17]–[19], [64]. However, lipid packing defects alone are not sufficient to induce membrane association, as OαS did not show any detectable interaction with zwitterionic DOPC vesicles. Notably, we have found that the secondary structure of free and membrane-bound OαS is very similar, indicating that a major structural rearrangement of these species does not take place upon membrane binding. This further suggests that the N-termini of the oligomer-forming monomeric subunits, which are crucial for oligomer anchoring into the bilayer, are folded in both free and membrane-bound OαS [59]. Nevertheless, OαS binding to the membranes occurs spontaneously under the experimental conditions, suggesting that oligomer-phospholipid interactions are more favourable than oligomer-solvent interactions, likely due to the high surface hydrophobicity of the amyloid oligomers in general [14], [64].

DOPS lipid membranes have previously been identified to strongly bind MαS and prevent the protein from aggregating [23]. However, here we show that MαS and OαS form a complex with liposomes of reduced electrophoretic mobility in comparison to free lipid vesicles, which corresponds to a lower ζ-potential. All particles investigated in this study presented ζ-potentials that indicate good to moderate stability in terms of classical colloidal science [54], [55].

While mounting evidence support the gain of function hypothesis of αS-mediated neurotoxicity in PD [13], [14], [65]–[67], very little evidence exists to support the loss of function hypothesis in PD ([68], reviewed in [69]–[71]). As the classic loss of function hypothesis in PD suggests, MαS is depleted from the functional MαS pool due to several reasons, including αS aggregation [69], [70]. Such MαS deficiency disrupts its physiological functions, including trafficking and maintaining the pool of synaptic vesicles or regulation of mitochondria numbers [70]. While minor MαS concentration shifts have not been shown to inhibit their function, when a significantly high fraction of MαS is deposited in the aggregates formed at later stages of the PD, the aforementioned downstream events become prevalent in the disease pathogenesis [72]. Here, we provide evidence of an alternative mechanism supporting the loss of function hypothesis that may play a role in the early stage of PD. While MαS and OαS coexist in the same cellular compartments, due to a 150-fold difference in their membrane binding affinities, OαS may outcompete and exclude MαS from the membrane surface. Hence, at the early stage of PD, the pool of non-aggregated MαS may still be available, but their access to the membrane is blocked. The MαS deficiency-related downstream effects can thus be initiated before a significant fraction of MαS is deposited in the inclusion bodies, compromising neurons healthy function much earlier than can be detected by measuring MαS concentration alone and accelerating the onset of PD [70].

In conclusion, capitalizing on an extensive biophysical characterisation of MαS- and OαS-lipid binding and their interplay on the membrane surfaces, we provide a new mechanism supporting the loss of function neurotoxicity hypothesis in PD. Not only does this work pave the way for further fundamental investigations on αS-linked PD mechanisms, but it also demonstrates how an array of powerful microfluidic methods enables in-depth insights into complex protein-lipid systems *in vitro*, which can further be expanded to the study of specific lipid membranes found at a range of cellular organelles. The platform presented here is not limited to αS and therefore has the potential provide insights into a wide range of amyloid disease mechanisms, the understanding of which is crucial for the development of effective therapeutics.

## Study Limitations

This study takes a systematic approach rooted in physical chemistry to delve into the interactions between MαS, OαS, and lipid bilayers. By employing a bottom-up methodology with simplified model systems for membranes and proteins, we gained precise control over experimental variables. This controlled approach is essential for unraveling the fundamental physical principles underlying the intricate binding energy landscape involved in Parkinson’s disease (PD) pathology.

Notably, our investigation concentrates exclusively on the binding energies of intrinsically disordered and misfolded alpha-synuclein, the central player in PD pathology. The study intentionally omits an in-depth exploration of structural aspects of alpha-synuclein-membrane binding. Instead, it is focused on the core binding energetics, offering a foundational understanding of MαS/OαS-lipid interactions.

Our findings serve as a pivotal cornerstone for comprehending the biophysical fundamentals of how MαS and OαS interact with lipid surfaces. Importantly, this work opens a new avenue for future research dedicated to understand the full molecular picture of PD mechanisms. Future investigations taking into account the complexities of cellular environments and additional molecular factors would offer a more comprehensive view of the roles of MαS and OαS in PD.

## Methods

### Materials

PDMS (polydimethylsiloxane) and curing agent were purchased from Dow Corning, PGMEA was obtained from Sigma-Aldrich, SU8-3050 from MicroChem Corporation. Carbon nanopowder was purchased from Fisher Scientific. 1,2-dioleoyl-sn-glycero-3-phospho-L-serine (DOPS) and 1,2-dioleoyl-sn-glycero-3-phosphocholine (DOPC) were obtained from Avanti Polar Lipids, DOPE-ATTO 647 from ATTO-Tec.

### MαS expression, purification and labelling

The N122C variant of αS was chosen as the amino acid at position no. 122 is not buried in the oligomer core and is not involved in αS binding to the membranes [13], [73]. Recombinant WT and N122C variants of αS were expressed in *E. coli* and purified as described [14]. To label the protein, the αS N122C variant was incubated with 1.5x molar excess of Alexa Fluor 488 C_5_-maleimide dye (ThermoFisher Scientific) in PBS at 4 °C overnight. The labelled protein was purified on a Superdex 200 16/300 column (GE Healthcare) to separate it from the unreacted dye. The concentration of labelled MαS was determined by measuring the absorbance of Alexa 488 dye at 495 nm (ε_495_ = 72 000 L·mol^-1^·cm^-1^) with a UV-VIS spectrophotometer. The labelled protein was aliquoted, flash-frozen in liquid nitrogen and stored at -80 °C until further use.

### Preparation of stabilised OαS

Stabilised OαS were prepared as described [14]. Briefly, 3–6 mg of lyophilized MαS was dissolved in 800 μl PBS, filtered with a 0.22 μm syringe filter and incubated overnight at 37 °C under quiescent conditions. The next morning, the samples were ultracentrifuged for 1 h at 90 000 rpm (288 000 g) at 20 °C to pellet amyloid fibrils. Ultracentrifugation was followed by filtration series with 100 kDa centrifugal filter units to remove MαS and small OαS. OαS were used within 2 days of preparation.

### Analytical ultracentrifugation

Sedimentation velocity analysis was carried out at 20 °C in a Beckmann-Coulter Optima XL-I analytical ultracentrifuge equipped with UV-visible absorbance optics and An50Ti rotor. The samples were centrifuged at 38 000-43 000 rpm (106 570-136 680 g) and monitored by the absorbance of the conjugated AlexaFluor-488 dye. The data were corrected to standard conditions by using the SEDNTERP programme. The distribution of sedimentation coefficients was determined by least squares boundary modelling of sedimentation velocity using the c(s) and ls-g*(s) methods implemented in the SEDFIT programme [74].

### Liposome preparation

Chloroform solutions containing desired lipids were mixed in glass vials. 1% DOPE-ATTO 647 was added to label the liposomes. Chloroform was then evaporated with a gentle nitrogen stream. The lipid film was hydrated with a 20 mM NaP buffer (pH 6.5 or 7.4) and vortexed. Lipid solutions were subjected to 5 freeze-thaw cycles in liquid nitrogen and a 42 °C water bath. To prepare SUVs, lipid solutions were sonicated on ice for 15 min using 50% cycling and 25% probe sonicator power. The residual titanium particles from the tip of the sonicator were spun down for 45 min at 15,000 rpm in a tabletop centrifuge at room temperature. The vesicles were then extruded via a 30 nm pore size membrane above the lipid phase transition temperature to obtain monodisperse liposomes, with a total of 31 passes. To prepare LUVs, the sonication step was omitted. Instead, lipids were passed through a 100 nm polycarbonate membrane, with a total number of passes being 31.

### Microfluidic diffusional sizing

Microfluidic diffusional sizing experiments were run on custom-made PDMS devices as described [22]. Before the experiments, the surface of microfluidic diffusional sizing devices was pre-treated with 0.01% Tween 20. The diffusional sizing experiments were run at 100–300 μl/h flow rates. Devices were equilibrated for 5 min before taking fluorescence images. To obtain the hydrodynamic radii values, the images were analysed with a custom-written Python script.

### Determination of dissociation constants and binding stoichiometries

The binding parameters for affinity and stoichiometry were determined using a two-state equilibrium model with no cooperativity. First, the apparent radii were converted to the molar fraction of αS bound to vesicles. For the oligomeric systems in which the fold-change in radii between free and bound αS is low, the relationship between fraction bound and apparent radius can be well approximated as linear. The fraction of OαS bound was therefore determined by the following equation:

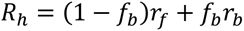

Where *R_h_* is the measured radius, *r_f_* and *r_b_* are the radii of the free and bound OαS, respectively, and *f_b_* is the fraction of OαS bound to vesicles. However, this linear approximation is not appropriate for the monomeric binding systems, where the relative difference between free and bound radii is larger. In these cases, a calibration curve was calculated by determining the apparent radii of linear combinations of diffusion profiles for free and fully bound MαS (Figure S3). These curves were approximated by second-order polynomials to convert apparent radii to the fraction of the bound monomer.

By solving the binding equilibrium equation, we obtain the following expression for the fraction of bound α-synuclein:

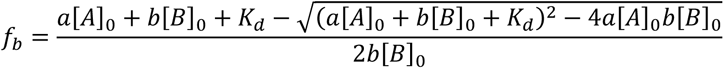

Where a is the effective number of α-synuclein particles bound per lipid, b is the number of monomers per particle, *[A]_0_* and *[B]_0_* are the total concentrations of lipid and α-synuclein, respectively, and is the dissociation constant in units of molar particle-lipid interactions. In the case of monomer binding, b is set to one. The posterior probability distributions for *K*_D_, *a*, and *b* (only in the case of OαS) were determined by Bayesian inference. The prior probability distributions were considered to be flat in logarithmic space for *K*_D_ and *a*, while sedimentation velocity analysis was used to determine an empirical size distribution (*b*) for the OαS. A normal distribution was used as the likelihood function, with the standard deviation as an inference parameter (prior flat in linear space) and mean defined by the above equation.

### Microfluidic free-flow electrophoresis devices fabrication

Microfluidic devices for µFFE were fabricated as described in [75], [76] and followed a similar procedure as the fabrication of MDS devices described above. Briefly, two masters with structures of the height of 50 µm were produced, one of them additionally containing 5 µm height channels for sample injection and junctions between electrolyte channels and the electrophoretic chamber. Upon casting and baking of clear PDMS, the individual devices from 2 masters were cut out, plasma treated for 30 s at 40% power and bonded together to produce a PDMS-PDMS device with a central electrophoresis chamber of 100 µm height and sample delivery port positioned in the middle of one of the chamber’s walls. Directly before the use, the devices were pre-treated with oxygen plasma for 800 s to activate the PDMS surface and filled with Milli-Q water to prolong the hydrophilicity of the surface.

### Microfluidic free-flow electrophoresis

The detailed operation of µFFE devices was described in Refs. [75], [76]. In brief, the performance of the electrophoresis relied on flowing the carrier medium (20 mM NaP, pH 6.5), sample and electrolyte from glass syringes (Hamilton) mounted on automated syringe pumps (NeMESYS, Cetoni GmbH) at 1000, 300 and 10 µl/h, respectively. The outlets of electrolyte solution (3 M KCl) were connected through metal pins to a source of voltage, thus creating liquid electrodes on the sides of the electrophoresis chamber. In the experiment, a voltage in increments of 20 V was applied from 0 to 80 V and fluorescent images of the sample stream deflected in an electric field were acquired. In parallel, readings of current were acquired. Each device was calibrated by replacing all liquids with electrolyte solution and measuring currents for determining the electrical resistance of the electrodes and estimating the effective electrical potential applied across the devices. The images were analysed with a custom-written Python script to construct electropherograms and obtain the values of electrophoretic mobilities.

### CD measurements

CD spectra in the 200–250 nm range were recorded with a Jasco far-UV CD spectrophotometer in a 1 mm path length cuvette. 5 μM of monomeric or OαS were mixed with 1 mM 110 nm DOPS LUVs in 20 mM NaP pH 7.4 buffer. The plots represent an average of 10 individual spectra of the same sample.

## Abbreviations

αS: α-synuclein
AD: Alzheimer’s disease
CD: Circular dichroism
DOPC: 1,2-dioleoyl-sn-glycero-3-phosphocholine
DOPE: 1,2-dioleoyl-sn-glycero-3-phosphoethanolamine
DOPS: 1,2-dioleoyl-sn-glycero-3-phospho-L-serine
LUV: Large unilamellar vesicle
MDS: Microfluidic diffusional sizing
µFFE: Microfluidic free-flow electrophoresis
MαS: Monomeric α-synuclein
OαS: Oligomeric α-synuclein
PD: Parkinson’s disease
PG: Phosphoglycerol
PS: Phosphoserine
PDMS: Polydimethylsiloxane
PGMEA: Propylene glycol methyl ether acetate
SUV: Small unilamellar vesicle

## Acknowledgements

We thank Timothy J. Welsh from Knowles Group for helpful discussion about the µFFE device preparation and data analysis. We thank Raphael Jacquät from the Knowles group for help with MDS data analysis. We thank Becky Gregory, Ewa Andrzejewska, Mollie McKeon and Ewa Klimont from the Centre for Misfolding Diseases for help with the purification of αS. This work was funded by the UK Engineering and Physical Sciences Research Council (EPSRC) grant EP/S023046/1 for the Centre for Doctoral Training in Sensor Technologies for a Healthy and Sustainable Future (G.Š.), Fluidic Analytics Ltd (G.Š. and M.A.C.), a Herchel Smith Research Studentship (C.K.X.), the Cambridge Trust (A.K.J.), the EPSRC grant EP/L015978/1 for the Centre for Doctoral Training for Nanoscience and Nanotechnology (NanoDTC) (A.K.J.), the Schmidt Science fellowship in partnership with the Rhodes Trust (K.L.S.) and St. John’s College Junior Research Fellowship (K.L.S.), the European Research Council under the European Union’s Horizon 2020 Framework Programme through the Marie Skłodowska-Curie grant MicroSPARK (agreement no. 841466; G.K.), and MicroProtLip (agreement no. 896068; M.A.C.), the Herchel Smith Fund of the University of Cambridge (G.K.), the Wolfson College Junior Research Fellowship (G.K.), the Ernest Oppenheimer Early Career Research Fellowship (A.L.). We would like to acknowledge funding from the European Research Council under the European Union’s Horizon 2020 research and innovation program through the ERC grant DiProPhys (agreement ID 101001615), the Frances and Augustus Newman Foundation (T.P.J.K.), the Medical Research Council (MRC) Career Development Award MR/W01632X/1 (J.R.K.) and the Centre for Misfolding Diseases (G.Š, A.K.J., M.A.C., C.K.X., A.L., G.K., T.P.J.K. and others).

## Author contributions

G.Š., M.A.C., C.K.X., A.K.J., G.K., S.D., H.F., J.R.K. A.L. and T.P.J.K. conceived the project. G.Š., M.A.C., C.K.X. and A.K.J. carried out the experiments and analysed the data. M.C.C. provided support with oligomer preparation. W.A. and T.M. provided support with instrumentation. Q.P., W.A. and K.L.S. helped with data analysis. All authors wrote and reviewed the paper.

