## Supplementary material for "α-synuclein oligomers displace monomeric α-synuclein from lipid membranes": SI information

### Supplementary figures

#### Characterisation of stabilised OαS

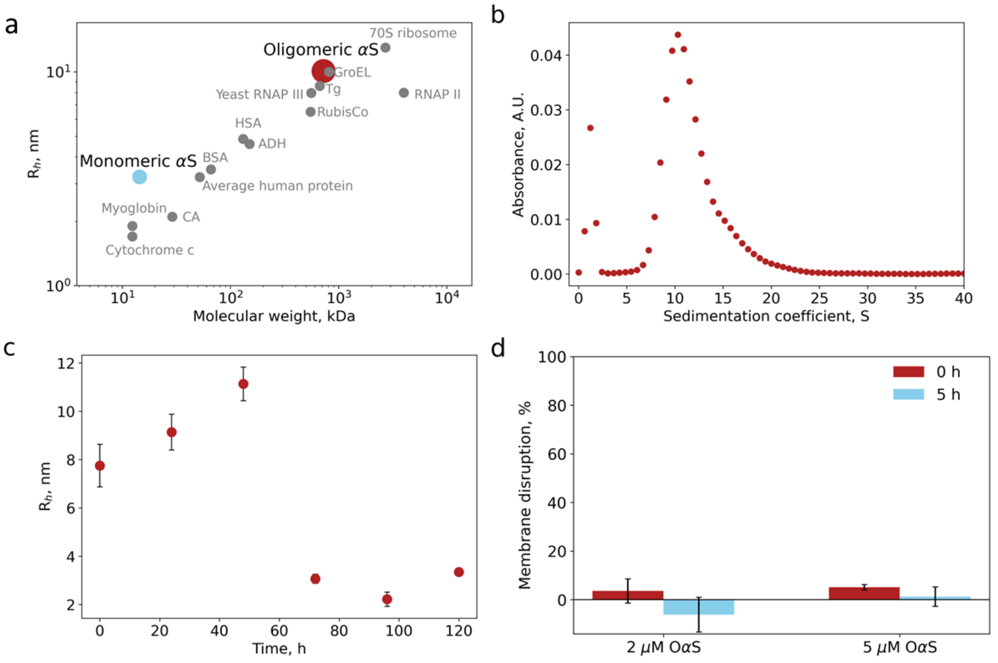


Figure S1. Characterisation of stabilised OαS. a) Hydrodynamic radii ($R_h$, nm) of MαS (blue) and OαS (red) αS as determined by microfluidic diffusional sizing and plotted as a function of molecular weight. Information about other proteins is retrieved from the database [1]. b) OαS size distributions as determined by analytical ultracentrifugation. c) OαS stability as determined by microfluidic diffusional sizing. OαS size distribution was monitored for up to 120 hrs, showing the limited stability of OαS prepared through lyophilisation. Error bars represent standard deviations of n = 3–4 measurements on individual microfluidic chips. d) OαS-induced membrane disruption kinetics, probed by the calcein dye leakage assay.

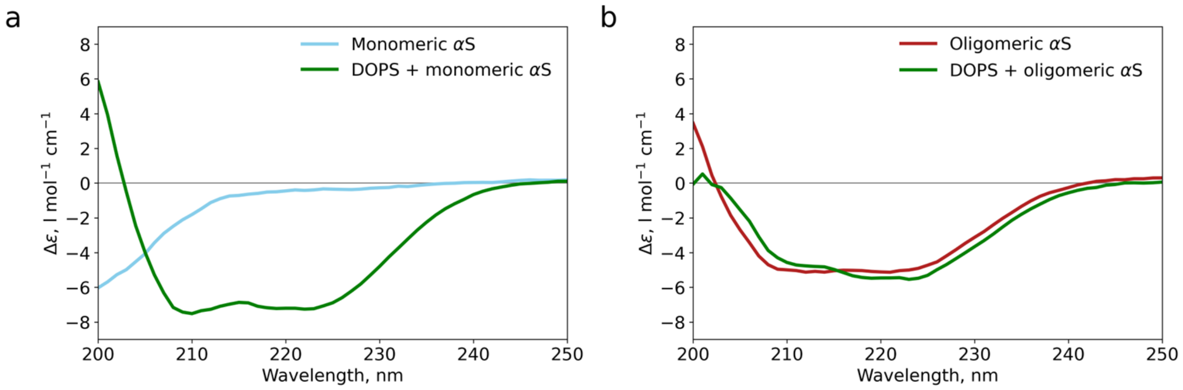


Figure S2. Background-subtracted CD spectra of free and DOPS LUVs bound (a) MαS and (b) OαS. 5 µM equivalent of MαS and 1 mM DOPS LUVs ($R_h$ = 60 nm) were used.

#### Calibration of diffusional sizing measurements

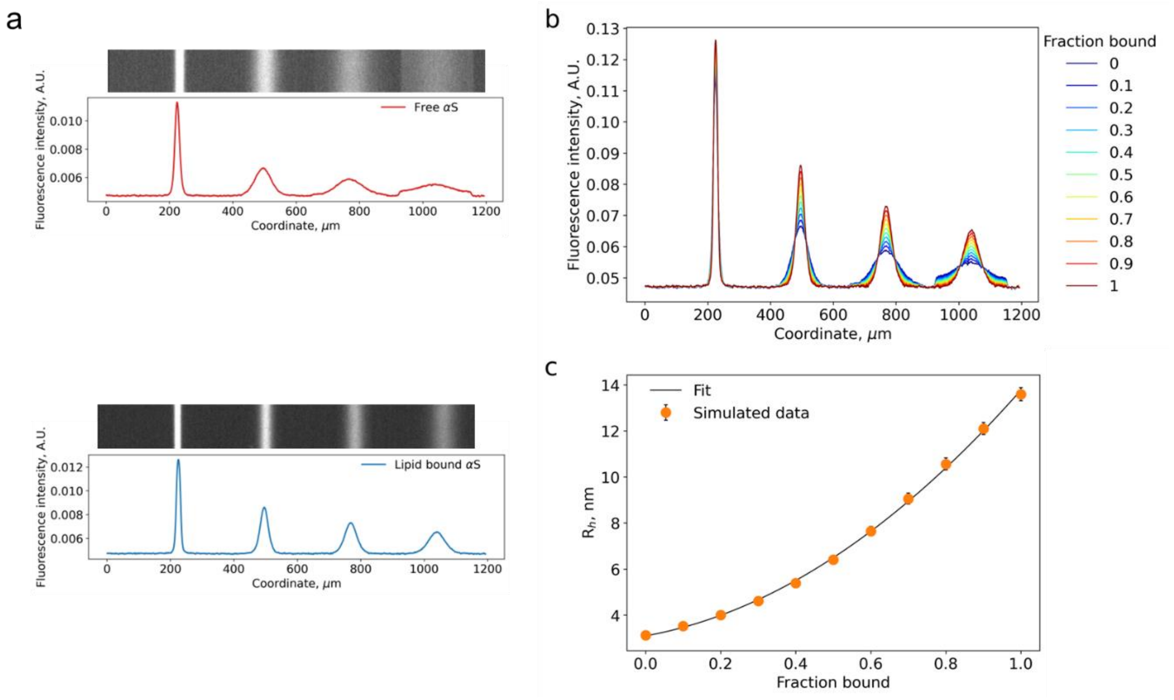


Figure S3. Calibration of diffusional sizing measurements. a) Diffusion profiles of MαS (top) and MαS-SUV complex (bottom). b) Linearly combined diffusion profiles of MαS and MαS-SUV complex at ratios 1:9 – 9:1. The simulated profiles correspond to the samples where a known fraction of MαS (from 0 to 1 or 0–100%) is bound to the DOPS SUVs. c) The calibration curve. $R_h$ plotted as a function of SUVs-bound fraction of MαS. 2 μM of MαS and 1 mM DOPS SUVs in 20 mM NaP pH 7.4 were used. Each peak corresponds to a different position in a microfluidic channel.

To accurately determine the bound fraction of αS using hydrodynamic radii as an input, we followed this procedure:

1. Diffusion profiles of free MαS were obtained by sizing 2 μM of MαS (Figure S4a, top).
2. Diffusion profiles of fully bound MαS were obtained. 2 μM of MαS were mixed with 1 mM DOPS SUVs, left to equilibrate at room temperature and then sized (Figure S4a, bottom).
3. Background-subtracted, area-normalized profiles obtained in steps 1 and 2 were then combined at various ratios to generate artificial diffusion profiles, each corresponding to a specific fraction bound value (from 0 to 1, incrementing fraction bound value by 0.1) (Figure S4b).
4. Simulated profiles were then fed into the custom-written MDS sizing script. Fraction bound values versus the obtained $R_h$ values were then plotted to obtain the calibration curve (Figure S4c).
5. The calibration data were then fitted to the polynomial function (2$^{nd}$ order polynomial normally).
6. The curve was then used to calculate the bound fraction of MαS using the measured radius as an input.

Calibration measurements were carried out in every set of affinity determination experiments (every liposome and protein prep). Diffusion profiles for calibration were obtained under as close conditions as possible (the same flowrate, same image exposure, devices of the same height, same αS concentration).

#### μFFE device design and operation

μFFE enables rapid in-solution analyte separation based on the differences in the electrophoretic mobility which is a function of the size and charge of molecules, hence it is suitable for addressing heterogenous samples (Figure S5) [3]–[7]. Deflection profiles acquired at varying voltages were analysed and electrophoretic mobility values for protein, lipid vesicles and protein-lipid complex were calculated. Generally, at a low DOPS:αS ratios, the signal observed in the protein channel (Alexa 488) separates into two streams, one of lower mobility, corresponding to the free protein and the other overlapping with the signal measured in the lipid channel (ATTO 647), corresponding to the lipid-protein complex (Figure S4b, bottom). At the highest ratio tested (500:1), the excess of LUVs separated from the stream of the αS-DOPS complex.

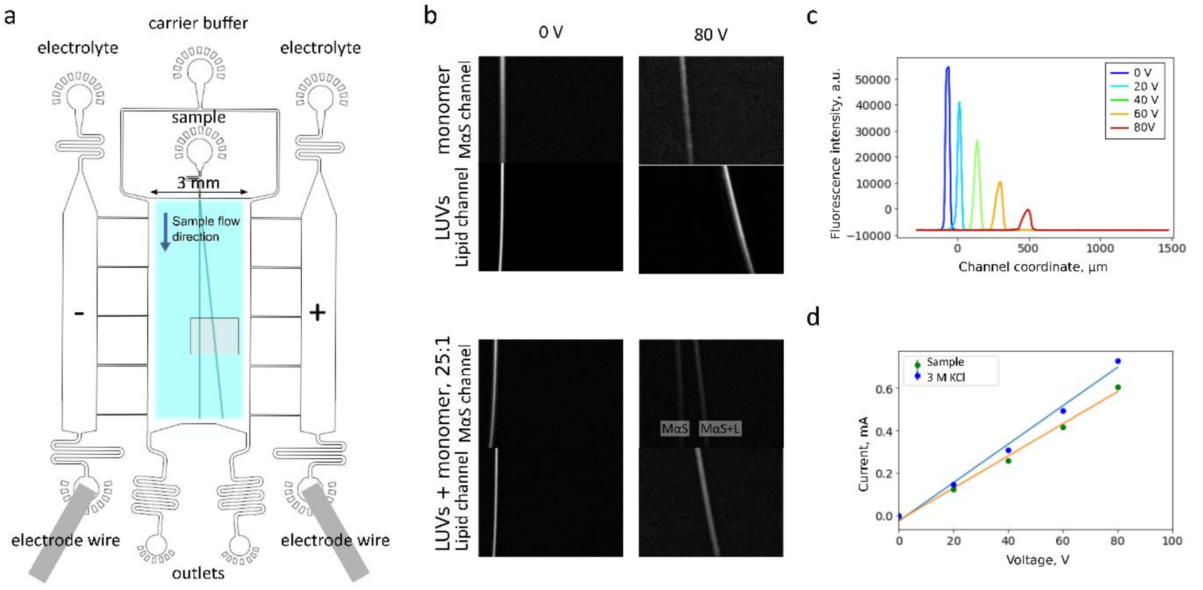


Figure S4. a) Scheme of the microfluidic device used for μFFE (top view). The shaded framed region indicates the image acquisition area. b) Exemplary images of samples in the electrophoretic chamberof pure LUVs/ MαS (top) and LUVs-MαS mixture at 25:1 lipid:protein ratio (bottom). c) Deflection profiles of an exemplary sample at different voltages used for electrophoretic mobility determination. d) Plot of the electrical current transmitted through the device both with the sample present (orange line) and when the device was filled with 3 M KCl electrolyte solution (blue line). The plot is used to determine the efficiency of each device.

#### Control experiments for the competition assay

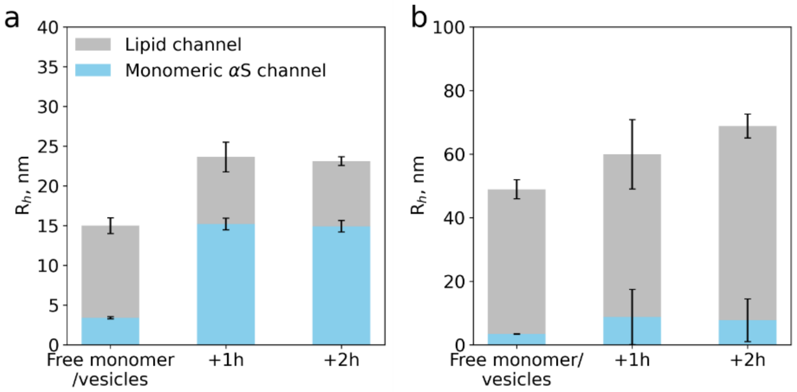


Figure S5. Data from an MDS control experiment demonstrating that MαS forms a complex with both SUVs (a) and LUVs (b) that is stable during the experimental timeframe (at least 2 hours). Error bars represent standard deviations of n = 3–4 measurements on individual microfluidic devices.

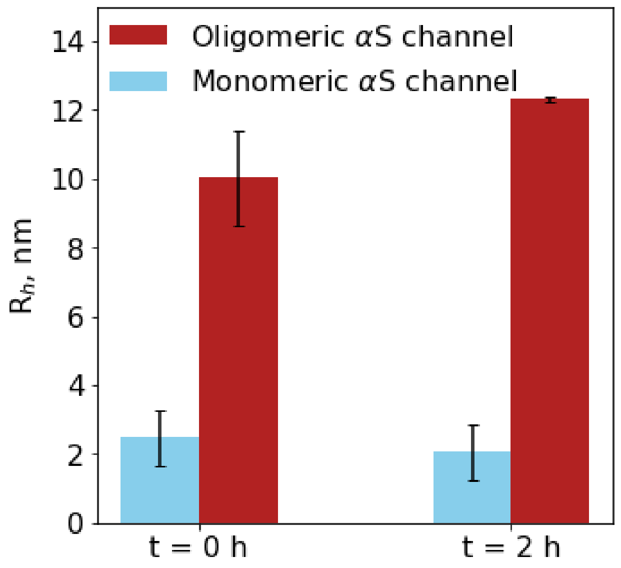


Figure S6. Size determination of co-incubated MαS and OαS with MDS. No change in size is observed for MαS and OαS indicating they co-exist in equilibrium within the experimental timescale. Error bars represent standard deviations of n = 3–4 measurements on individual microfluidic devices.

Table S1. Best fit values for dissociation constants and binding stoichiometries of MαS/OαS-membrane binding. 95% credible intervals are given in square brackets for those which yielded well constrained distributions.

| Sample | Dissociation constant ($K_D$) (This study) | Stoichiometry (effective lipid molecules per MαS or OαS) (This study) | Dissociation constant ($K_D$) (Literature) | Stoichiometry (Literature) |
|---|---|---|---|---|
| MαS-SUVs 6.5 | 0.077 μM [0.018, 0.18] | 23 [17, 28] | 0.2 ± 0.1 μM [8]<br>0.29 ± 0.25 μM [9]<br>0.05 μM [9] | 33 ± 1 [8]<br>57.2 ± 4.0 [9]<br>78 [9] |
| OαS-SUVs 6.5 | <0.6 μM | 658 [23, 2566] | N/A | N/A |
| OαS-LUVs 6.5 | >2 μM | 3<3000 | N/A | N/A |
| MαS-SUVs 7.4 | <30 μM | 21 [2, 35] | N/A | N/A |
| OαS-SUVs 7.4 | 3.5 nM | <2311 | N/A | N/A |
| OαS-LUVs 7.4 | 21 nM [2.5, 1100] | 4809 [152, 20820] | N/A | N/A |
| MαS-SUVs 7.4 High salt | >1 μM | <208 | N/A | N/A |

### Supplementary Methods

**Preparation of calcein-loaded DOPS liposomes**

First, 0.6 g calcein was dispersed in 4 ml MilliQ water followed by addition of 6 M NaOH while stirring vigorously. Calcein concentration was then confirmed by a UV-Vis spectrophotometry by measuring the absorbance at 490 nm ε = 72 000 $L \cdot mol^{-1} \cdot cm^{-1}$. To prepare calcein-encapsulated LUVs, the formation of a dry lipid film was followed by lipid dispersion in calcein solution. Lipids were then extruded via a 100 nm polycarbonate filter (number of passes 31). Lipid vesicles were then purified into the assay buffer (20 mM Hepes, 120 mM NaCl 0.8 μM EDTA pH 8.0) on PD-10 columns loaded with Sephadex G-50. Vesicle size was then determined by DLS.

**Membrane permeabilization (calcein release) assay**

Samples containing calcein-loaded 50 μM DOPS liposomes were mixed with 2 μM and 5 μM OαS and aliquoted to 96-well plates (Corning 3651). The plate was subjected to 300 rpm double orbital shaking at 37°C with simultaneous fluorescence readout from the top at 488 nm excitation and 520 nm emission in a microplate reader. The fluorescence values were normalised taking into account the vesicle background fluorescence data and maximum dye release fluorescence intensity values by using the equation:

$$Membrane\ disruption, \% = \frac{I - I_{min}}{I_{max} - I_{min}}$$

Where I - fluorescence intensity at time t, $I_{min}$ fluorescence intensity background (vesicles in a buffer), $I_{max}$ - maximum fluorescence intensity signal (100% calcein release, vesicle sample with 5% Triton X-100).

#### Zeta potential determination

**Zeta potential.** The Henry equation was used to calculate zeta potential:

$$\zeta = \frac{3\mu\eta}{2\varepsilon_0\varepsilon_r f_H(\kappa R_h)}$$

Where $\varepsilon_0$ is the vacuum permittivity, $\varepsilon_r$ is the solvent dielectric constant, $\eta$ is the solvent viscosity and $f_H$ is the Henry function, approximated as:

$$f_H(\kappa R_h) = 1 + \frac{1}{2(1+\delta)^3},$$

$$\delta = \frac{5}{2\kappa R_h(1+2e^{-\kappa R_h})}.$$

Average values of radii ($R_h$) were used: MαS 2.57 nm, OαS 8.50 nm, MαS + LUVs 76.20 nm, MαS + SUVs 10.79 nm, OαS + LUVs 52.44 nm, OαS + SUVs 14.85 nm, LUVs 67.76 nm, SUVs 10.31 nm.
